# Inactivated rabies virus vectored SARS-CoV-2 vaccine prevents disease in a Syrian hamster model

**DOI:** 10.1101/2021.01.19.427373

**Authors:** Drishya Kurup, Delphine C. Malherbe, Christoph Wirblich, Rachael Lambert, Adam J. Ronk, Leila Zabihi Diba, Alexander Bukreyev, Matthias J. Schnell

## Abstract

Severe acute respiratory syndrome coronavirus 2 (SARS-CoV-2) is an emergent coronavirus that has caused a worldwide pandemic. Although human disease is often asymptomatic, some develop severe illnesses such as pneumonia, respiratory failure, and death. There is an urgent need for a vaccine to prevent its rapid spread as asymptomatic infections accounting for up to 40% of transmission events. Here we further evaluated an inactivated rabies vectored SARS-CoV-2 S1 vaccine CORAVAX in a Syrian hamster model. CORAVAX adjuvanted with MPLA-AddaVax, a TRL4 agonist, induced high levels of neutralizing antibodies and generated a strong Th1-biased immune response. Vaccinated hamsters were protected from weight loss and viral replication in the lungs and nasal turbinates three days after challenge with SARS-CoV-2. CORAVAX also prevented lung disease, as indicated by the significant reduction in lung pathology. This study highlights CORAVAX as a safe, immunogenic, and efficacious vaccine that warrants further assessment in human trials.

## Introduction

More than 150 vaccines against SARS-CoV-2 are in preclinical trials, and over 51 candidate vaccines are in human trials (1). They include adenovirus and other viral vector-based vaccines and protein-, DNA-, and mRNA-based vaccines (2).

These vaccine approaches have different advantages and disadvantages. mRNA vaccines, including the 2 U.S. FDA-approved coronavirus vaccines from Moderna and Pfizer-BioNTech, can be produced efficiently, but they can be costly to produce and have temperature sensitivity (3); furthermore, questions remain unanswered about the longevity of the immune response with mRNA vaccines and if they block transmission. Meanwhile, DNA-based vaccines benefit from temperature stability and low production costs, but their immunogenicity has been a concern, and the need for multiple vaccinations challenges their feasibility (4). Virus-like particle (VLP)-and recombinant protein-based vaccines have an excellent safety advantage because they do not replicate in the host, and often they can also be made more temperature stable. However, VLP-based vaccines are not always as immunogenic as replication-competent or replication-deficient viral vector vaccines and often require an adjuvant to increase their immunogenicity to an adequately protective level (5, 6). Finally, viral vector-based vaccines the advantage of being typically cheaper to produce, efficacious after a single vaccination, and often highly immunogenic (1, 7-9), but those based on live viral vaccines sometimes fail because of safety concerns and production scalability, and they usually require a low storage temperature (8). The disadvantages of viral vector vaccines are largely overcome, however, when they are based on an inactivated virus.

Using this approach, we have developed an inactivated viral vector vaccine against SARS-CoV-2 that is based on the rabies virus (RABV). We have previously utilized this approach to develop inactivated RABV-based vaccines for several other human pathogens (e.g., Ebola virus, Marburg virus, Lassa Fever virus and others) (10-27). These rabies virus-based vaccines have been proven to be highly immunogenic and protective against several emerging viral infections and bacterial toxins, as well as safe and temperature stable (28). The RABV vaccine itself has been safely used for decades in more than 100 million people, including children, pregnant women, and the elderly, and proven to result in long-lasting immunity (29).

CORAVAX, our rabies vectored SARS COV-2 vaccine, expresses the S1 domain of the SARS-CoV-2 spike (S) protein fused to part of the N terminal domain of the RABV glycoprotein (G) and is incorporated in RABV particles. Mice immunized with adjuvanted CORAVAX developed potent neutralizing antibodies about 5-10 times higher than the virus-neutralizing antibodies (VNA) detected in convalescent sera from SARS-CoV-2 infected people (30). Here, to learn more, we evaluated the efficacy of CORAVAX in the Syrian hamster model for SARS-CoV-2, which is currently considered the best model to study COVID-19 therapies and therapeutics. Our results showed that two immunizations with the adjuvanted CORAVAX vaccine induced high VNA and cleared infectious SARS-CoV-2 from the lung and the nasal turbinates on day 3 post challenge. Moreover, the pathology induced by SARS-CoV-2 was reduced with only some early infiltration of cells into the lung tissue. Taken together, this study indicates that CORAVAX warrants further development as a disease-preventing human vaccine.

## Results

### Immunogenicity of CORAVAX in Syrian hamsters

CORAVAX a rabies vectored SARS COV-2 vaccine was generated by inserting the S1 subunit of the SARS-CoV-2 Spike into the SAD-B19 rabies vaccine strain. To promote the incorporation of the S1 domain, we prepared a fusion protein between SARS-CoV-2 S1 and RABV G. Toward this approach, the N-terminal 682 aa of SARS-CoV-2 S1 were fused to a truncated RABV glycoprotein, which comprises 31 aa of the ectodomain (ED) of RABV G and the complete cytoplasmic domain (CD) and transmembrane domain of RABV G to allow chimeric glycoprotein incorporation into RABV virions. We previously showed that CORAVAX induces high virus neutralizing antibodies as a live or inactivated vaccine in mice (30). Here we extend these studies in the more relevant SARS CoV-2 Syrian Hamster model of severe disease.

To evaluate the immunogenicity and efficacy of the CORAVAX vaccine in golden Syrian hamsters, we vaccinated each group comprising 12 animals with either 10 μg of inactivated CORAVAX or control vaccine FILORAB1 (rabies vectored Ebola vaccine) adjuvanted with MPLA-AddaVax in a 100 μL injection volume via the intramuscular route (Figure 1A). The animals received a prime on day 0 and a booster on day 28. Blood was collected on days -2, 26, and 56 (Figure 1B). On day 60, vaccinated and control hamsters were challenged intranasally with a dose of 10^5^ PFU of the SARS-CoV-2 isolate USA_WA1/2020 (31).

**Figure 1:**
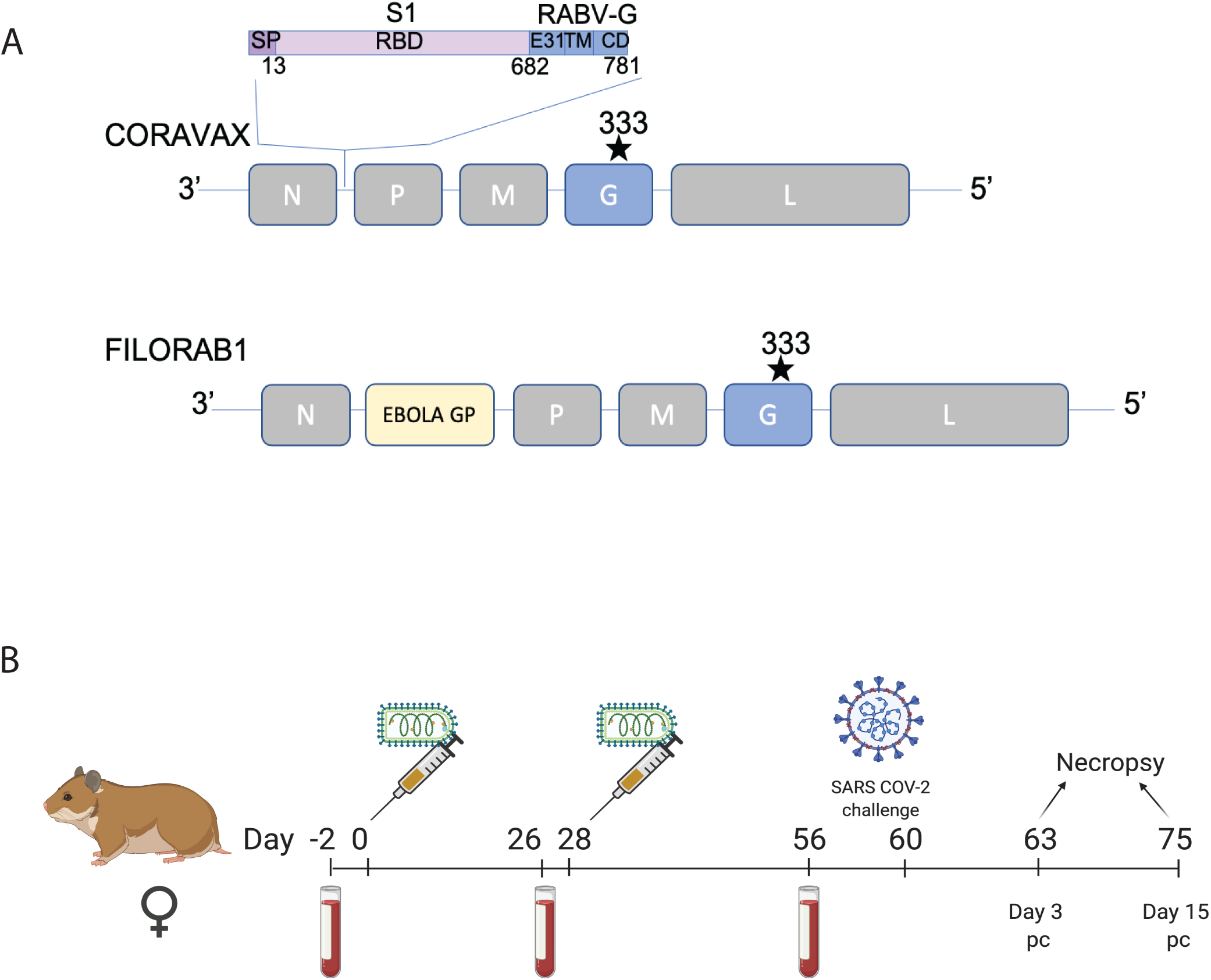
Vaccine constructs and vaccination schedule. A) Schematic illustration of CORAVAX, the rabies virus-based SARS-CoV-2 vaccine construct wherein a SARS-CoV-2 S1 RABV G chimeric protein cDNA was inserted between the N and P genes of the SAD-B19-derived RABV virus vaccine vector BNSP333 and control vaccine FILORAB1, the rabies virus-based EBOLA vaccine used in this study. B) Syrian hamsters were immunized on day 0 and day 28 with 10 μg chemically inactivated CORAVAX or FILORAB1 with MPLA-AddaVax adjuvant. The animals were challenged on day 60 with a dose of 10^5^ PFU of the SARS-CoV-2 isolate USA_WA1/2020. Serum was collected from each hamster at days -2, 26, 56, 63, and 75 for analysis. Animals were necropsied at days 63 (day 3 p.c.) and 75 (day 15 p.c.).

In the next step, sera from immunized hamsters and controls were assayed for SARS CoV-2 specific antibody responses by ELISA specific for SARS-CoV-2 S1 and receptor binding site (RBD) (Figure 2). High titers of SARS-COV-2 S1 specific IgG responses were detected in the CORAVAX vaccinated hamsters on day 26 with no significant differences in antibody titers on day 56 (Figure 2A). As expected, no SARS-CoV-2 S1 immune response was detected in the control animals vaccinated with the EBOV vaccine FILORAB1. As previously seen for other inactivated and adjuvanted RABV-based vaccines, CORAVAX induced a Th1 biased immune response as indicated by the high SARS CoV-2 S1 IgG2/3 responses detected on days 26 and 56 (Figure 2B). We could not detect S1 specific IgG1 responses in any of the hamsters (data not shown). Similar to the responses detected by ELISA to SARS CoV-2 S1, we observed high titers of SARS-CoV-2 VNA with mean titers of 976 ± SD 685 on day 56 (Figure 2C). Of note, no VNA against SARS-CoV-2 were detected in FILORAB1 immunized animals before the challenge (Figure 2C).

**Figure 2.**
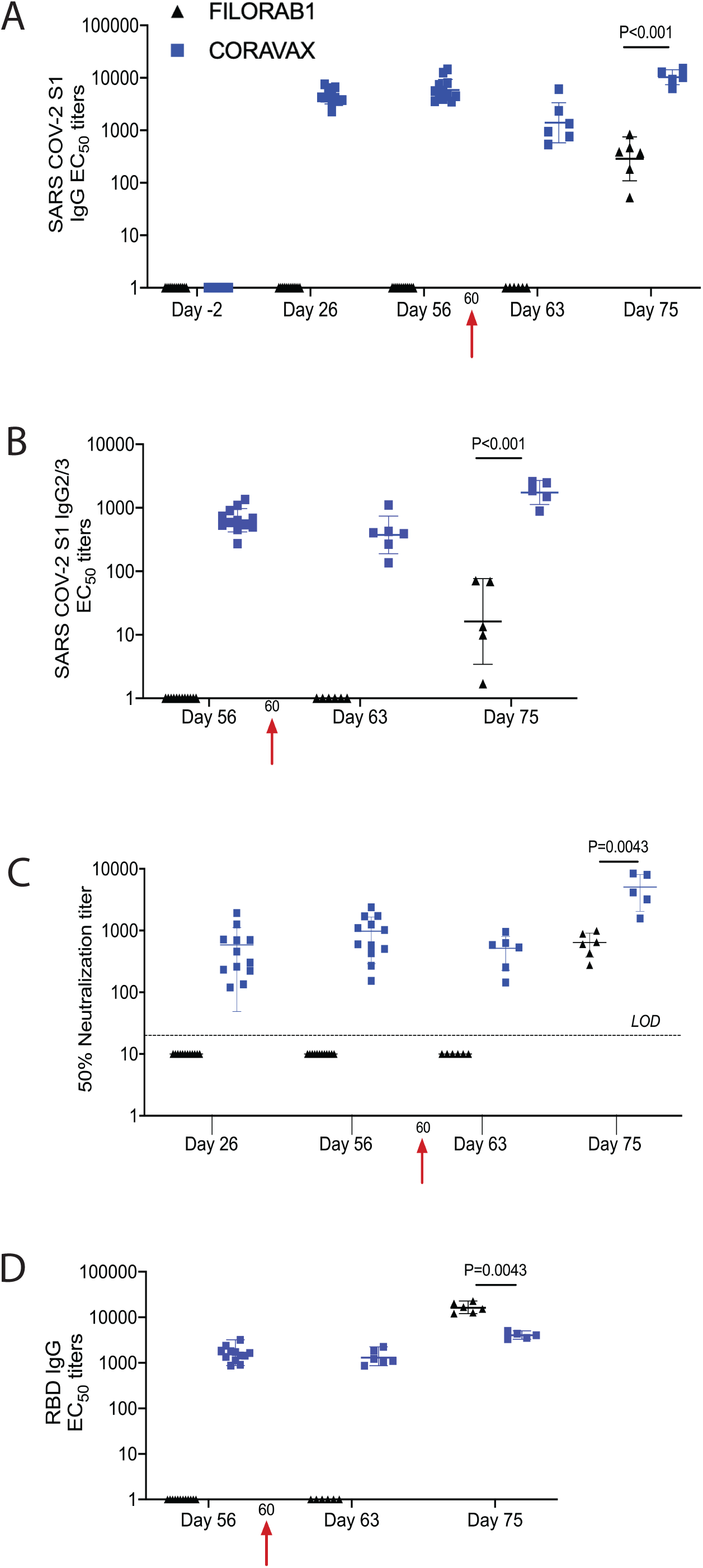
SARS CoV-2 immune responses. Serum samples collected from each hamster were evaluated for SARS-CoV-2 S-specific immune responses by A) ELISA, Anti-SARS CoV-2 S1 IgG responses represented as EC50 titers over time, B) ELISA, Anti-SARS CoV-2 IgG2/3 responses, C) Virus neutralizing antibodies, and D) ELISA, Anti-SARS COV-2 S RBD IgG responses. The CORAVAX vaccine group is shown in blue and the FILORAB1 group in black. For A-D, mean titers ± SD are depicted for each group per time point. P values determined by Mann-Whitney test. Only significant differences are depicted.

We next analyzed the immune responses post-challenge (p.c.). By day 63, 3 days p.c., we observed a significant reduction (P=0.0069) in S1 IgG responses in CORAVAX vaccinated animals, while no S1-specific IgG responses were observed in the controls hamsters (FILORAB1) (Figure 2A,C). By day 75, both in CORAVAX and FILORAB1 vaccinated and challenged hamsters, S1-specific IgG and VNA were detected (Figure 2A,B,C). However, both S1-specific IgG and VNA were significantly higher in the CORAVAX vaccinated animals on day 75 (Figure 2A, C). Moreover, p.c. CORAVAX vaccinated animals indicated a more robust Th1-biased immune response compared to FILORAB1 control animals, as indicated by the IgG2/3 responses (Figure 2B). In addition to the antibodies directed against the receptor-binding domain (RBD) SARS-CoV-2, high RBD-specific IgG titers were detected in the CORAVAX vaccinated animals on day 56 as well as p.c. on days 63 and 75. Interestingly, the RBD IgG titers were undetectable in the FILORAB1 vaccinated controls on day 63 but were higher than the ELISA titers detected in CORAVAX vaccinated animals on day 75, challenging the dogma that high ELISA titer against RBD predict the VNA against SARS-CoV-2 (Figure 2D).

#### CORAVAX induces potent immune responses against RABV

We also analyzed the VNA induced by the two vaccines against RABV. CORAVAX is a vaccine against both SARS-CoV-2 and RABV, and in large part of the world RABV is still a significant problem, <<annually >> killing about 55,000 people, mostly children. We detected a titer of anti-RABV neutralizing antibodies above the WHO’s 0.5 IU standard in both the FILORAB1 and CORAVAX vaccinated animals, with no significant differences (Figure 3).

**Figure 3.**
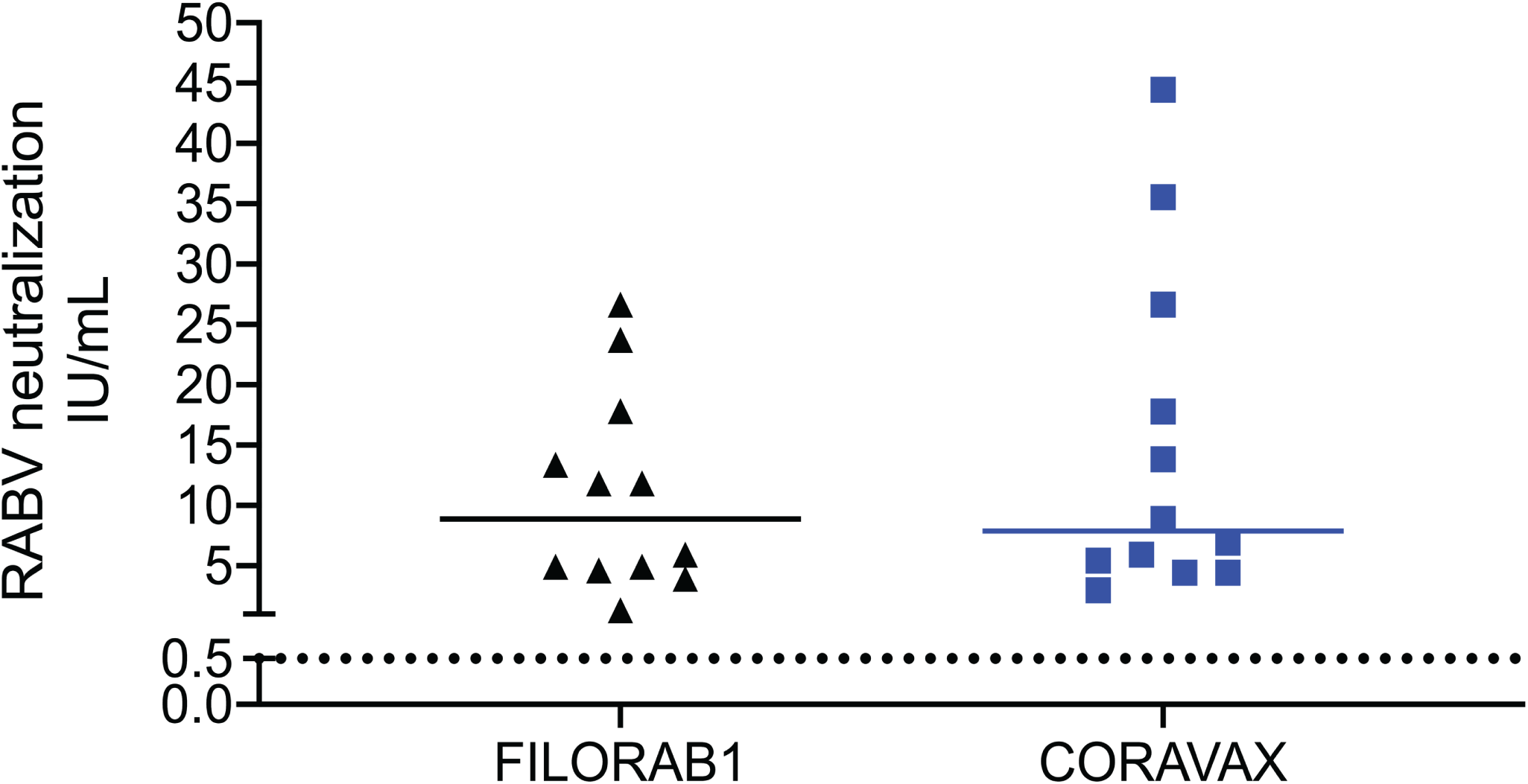
Rabies virus neutralizing antibodies. Hamster day 56 sera were assessed for RABV VNA. Dotted line represent 0.5 I.U/mL. Only significant differences are depicted.

#### CORAVAX protects the hamsters from weight loss and viral burden in the lungs and nasal turbinates post-SARS CoV-2 challenge

The hamsters were challenged intranasally on day 60 with 10^5^ PFU of SARS-CoV-2 isolate USA_WA1/2020 and were monitored for up to 15 days. The CORAVAX vaccinated animals showed significantly less weight loss than the FILORAB1 controls (P=0.0098), which lost more than 10% weight and recovered only at day 11 (Figure 4). At days 3 and 15 p.c., half of the hamsters in each study group were euthanized, and lungs and nasal turbinates were harvested and virus isolated to determine viral loads by plaque reduction assay (Figure 5A, B). Additionally, the number of viral copies was analyzed by RT-qPCR assay (Figure 5C, D). The lack of weight loss coincided with the absence of any infectious virus in the lungs and nasal turbinates of the CORAVAX vaccinated animals on days 3 and 15 p.c. (Figure 5A and B).

**Figure 4.**
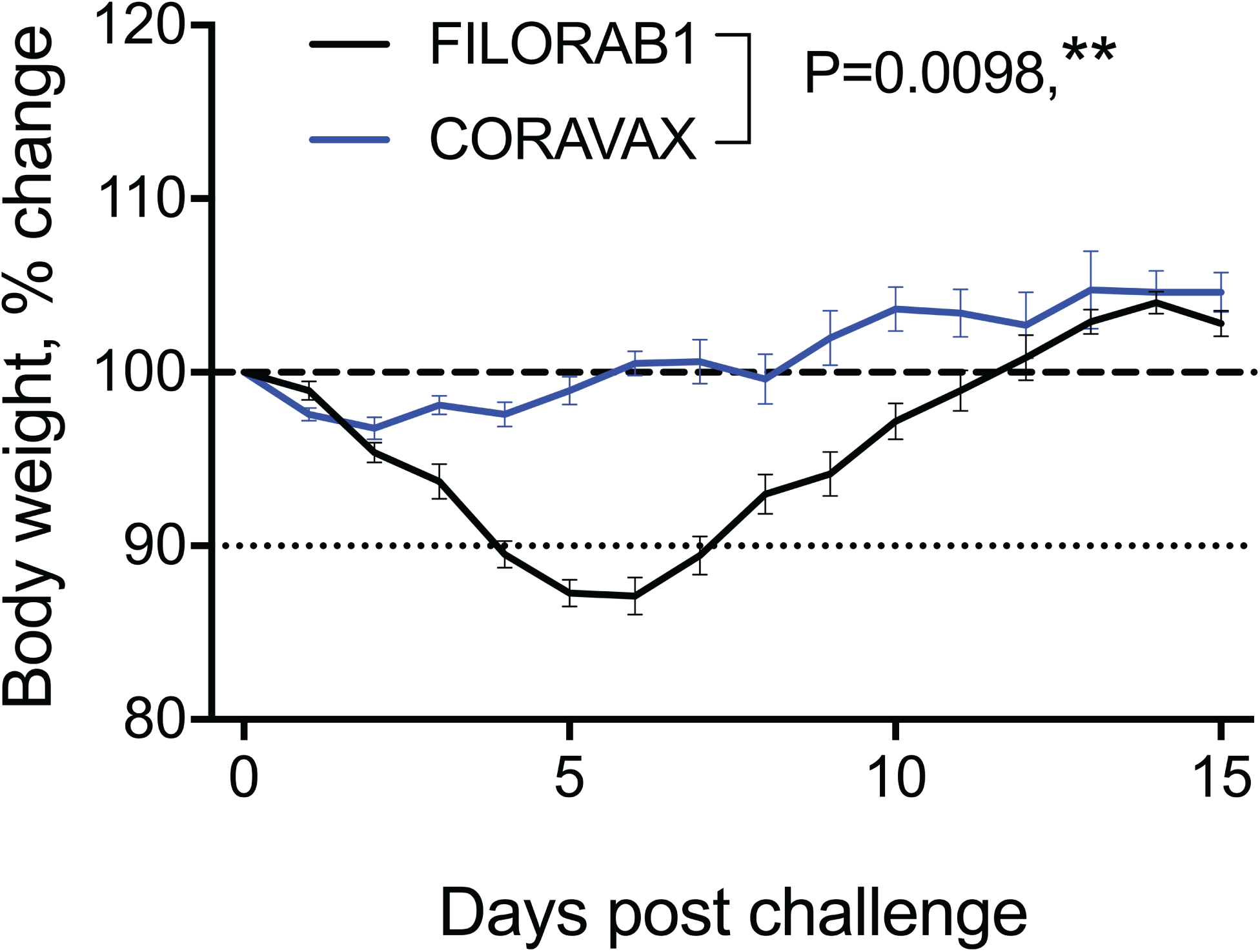
Hamster body weight after SARS CoV-2 infection. Hamsters were vaccinated at day 0 and day 28, challenged intranasally with 10^5^ PFU SARS-CoV-2 at day 60. Percent change in body weight. CORAVAX vaccine group is shown in blue and FILORAB1 group in black. N = 12 for FILORAB1 group (6 hamsters euthanized at day 3 p.c.) and N = 11 for CORAVAX group (6 hamsters euthanized at day 3 p.c.). Body weight P value determined by Wilcoxon test. P > 0.123 (ns), P < 0.033 (*), P < 0.002 (**), P < 0.001 (***).

**Figure 5.**
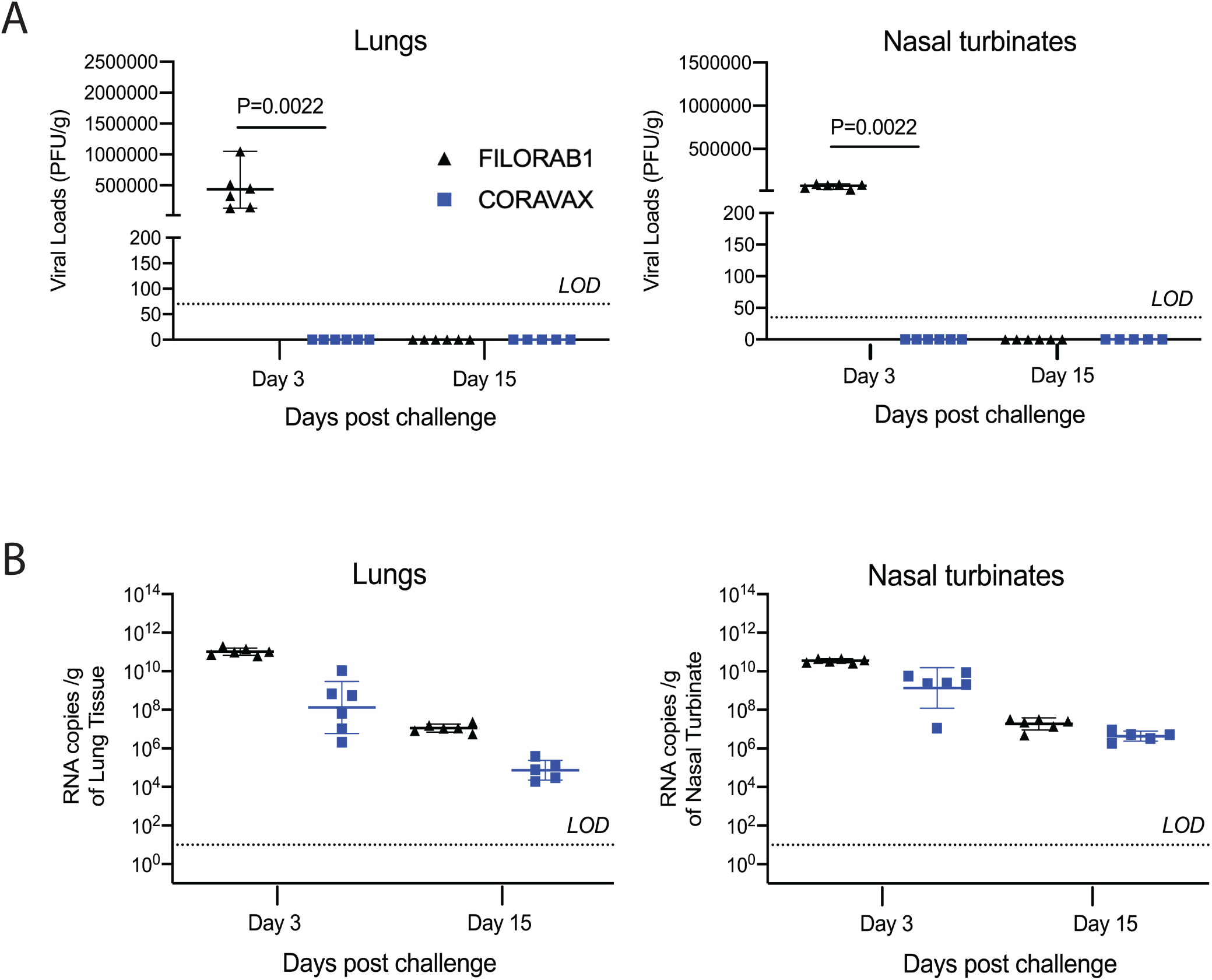
SARS-CoV-2 tissue viral load in hamsters. Hamsters were challenged intranasally with 10^5^ PFU SARS-CoV-2, and half of the animals in each group were euthanized at days 3 and 15 p.c. Right lungs (A, C) and nasal turbinates (B, D) from each animal were homogenized in media and viral loads were determined by plaque assays on Vero E6 cells (A, B) or by qRT-PCR (C, D). The limit of detection for the plaque assay was 70 PFU per lung and 35 PFU per nasal turbinate. The limit of detection for the qRT-PCR assay is 10 copies. The CORAVAX vaccine group is shown in blue and the FILORAB1 group in black. Data represent mean ± S.D., N = 6 for FILORAB1 group day 3 and day 15 time points, and N = 6 for CORAVAX group day 3 and N = 5 for CORAVAX group day 15 time points. P values determined by Mann-Whitney test. Only significant differences are depicted.

In contrast to the CORAVAX immunized animals, the FILORAB1 controls had high titers of infectious virus in the lungs and nasal turbinates on day 3 p.c.. As expected, both groups cleared the SARS-CoV-2 15 days.

A similar trend between the two groups was detected when RNA copies of SARS-CoV-2 were analyzed via RT-PCR. CORAVAX vaccinated animals had significantly reduced RNA copies in the lungs and nasal turbinates at necropsy (day 3 and 15 p.c.) than the control FILORAB1 group (Figure 5C, D). The presence of viral RNA in the absence of infectious SARS-CoV-2 is well-established and based on the stability of the genome of SARS-CoV-2 (32, 33). However, it should be noted that the ∼1000-fold lower copy number in CORAVAX vaccinated animals indicates a significant reduction of viral replication by CORAVAX.

#### CORAVAX vaccinated animals have significantly reduced lung pathology than controls

Lung sections were collected from control and vaccinated animals at days 3 and 15 p.c. (Figure 6, 7). Sections were scored in a blinded manner. Histopathological changes consistent with viral interstitial pneumonia were noted in all animals, regardless of treatment or time of collection (Figure 6, representative pathology pictures; Figure 7A, B mean overall pathology scores). These included consolidation, widespread alveolar septal thickening, and airway pathology consisting of airway epithelial hyperplasia and accumulation of inflammatory cells in airways, occasionally leading to obstruction of the lumen. On day 3 p.c., CORAVAX vaccinated animals had significantly lower average pathology scores. Specifically, component scores for inflammatory foci size and number and airway pathology were improved (Figure 7A). Animals cleared the virus by day 15 (Figure 5A, B), and consistent with expected tissue damage repair following clearance, we observed reduced pathology in both CORAVAX and the control FILORAB1 vaccinated animals (Figure 7B).

**Figure 6.**
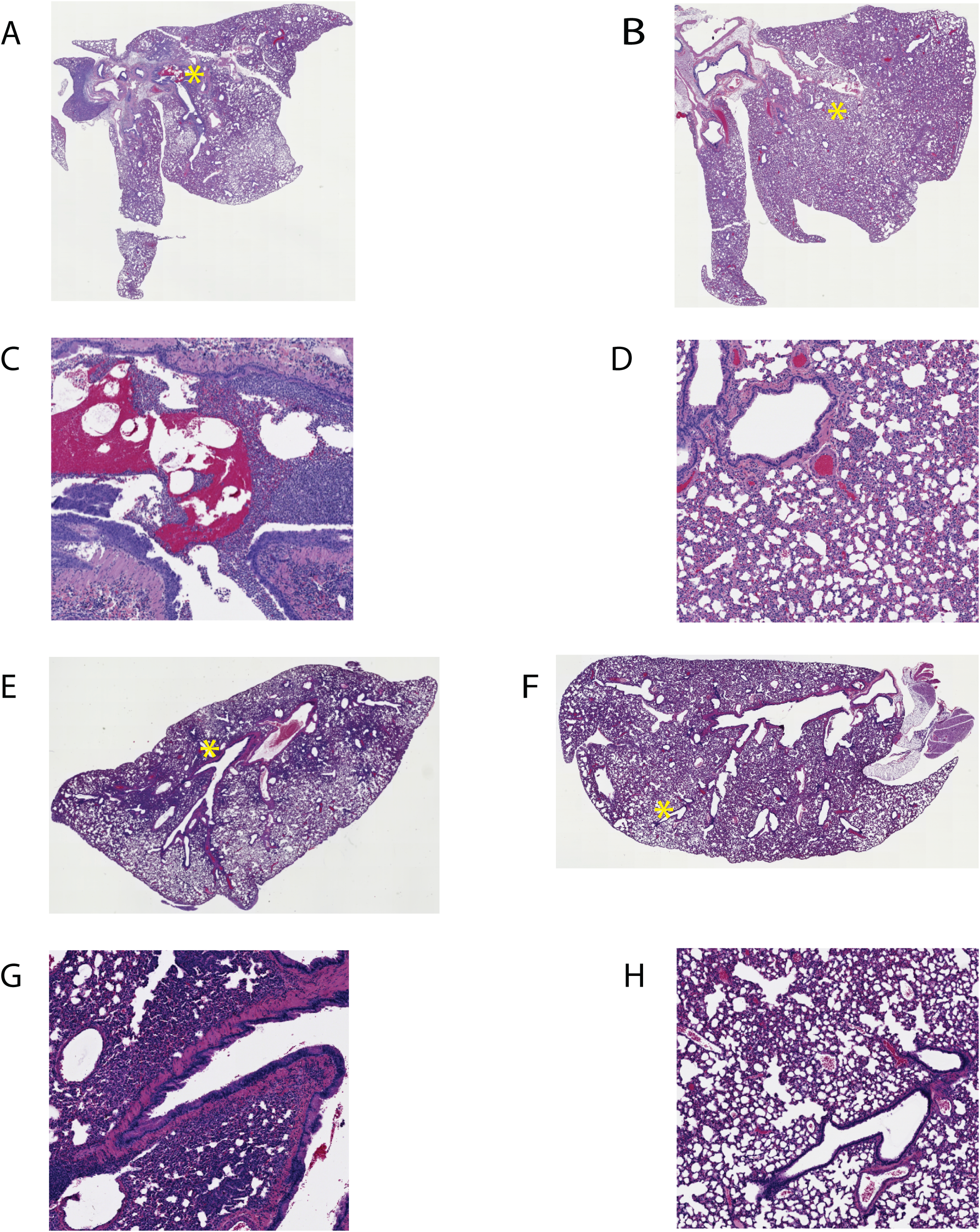
SARS-CoV 2 lung pathology. Representative histological images of SARS-CoV-2 infection in control and vaccinated hamster lungs. A and B, low magnification images, day 3 p.c. lungs. A: FILORAB1 control, B: CORAVAX vaccinated. C and D, high magnification of A and B detailing specific pathological features, location indicated by asterisks on low magnification images. Infiltration of parenchyma and airways by mononuclear inflammatory cells is prominent in control animal. Notable improvement in airway infiltration observed in vaccinated animals. Varying degrees of consolidation and septal thickening are present in control and vaccinated animals. E and F, low magnification images, day 15 p.c. lungs. E: FILORAB1 control, F: CORAVAX vaccinated. G and H, high magnification of E and F detailing specific pathological features, location indicated by asterisks (*) on low magnification images. Significant airway infiltration is absent in all animals, but other aspects of inflammatory pathology persist, such as epithelial hyperplasia Overall consolidation improved in vaccinated animals but did not reach statistical significance

**Figure 7.**
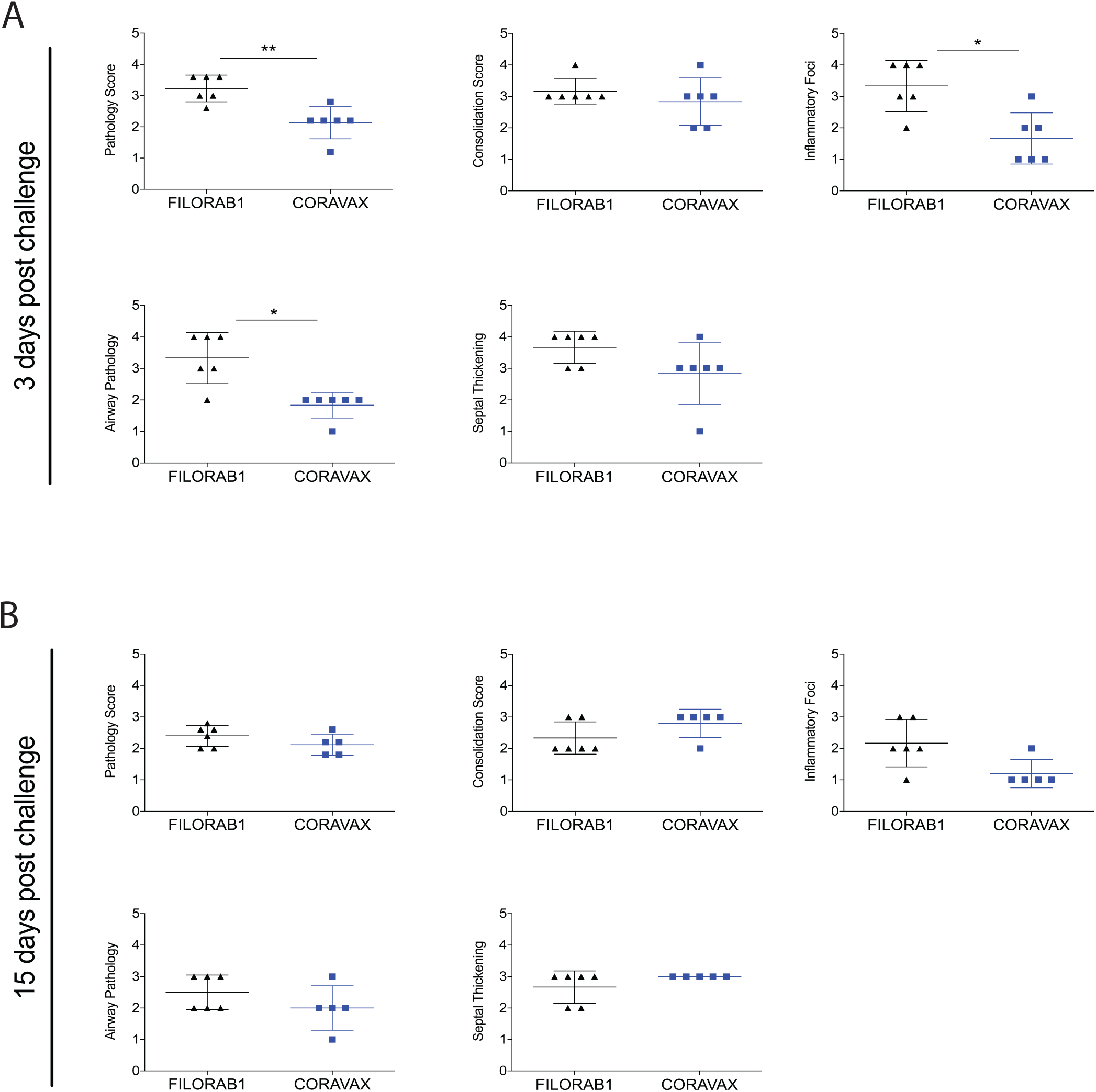
Comparative pathology scores for lungs from CORAVAX vaccinated and control hamsters post SARS-CoV-2 challenge. Scores at 3 days (A) and 15 days (B) p.c. Scores are displayed for overall lung pathology and individual criteria, including consolidation or extent of inflammation, type inflammatory foci, airway pathology, and septal thickening. The pathology scores (mean) were calculated based on the criteria described in Supplemental Table 1. The CORAVAX vaccine group is shown in blue and the control group in black. Data represent mean ± S.D., N = 6 for FILORAB1 group day 3 and day 15 time points and N = 6 for CORAVAX vaccine group at day 3 and N=5 for CORAVAX day 15 time points. P > 0.123 (ns), P < 0.033 (*), P < 0.002 (**), P < 0.001 (***). Only significant differences are depicted.

## Discussion

There is an urgent need for a safe and effective vaccine against SARS-COV-2 that can be administered to pregnant women, children, elderly and the immunocompromised. The two FDA emergency use authorization (EUA) COVID-19 vaccines, BNT162b2 (Pfizer-BioNTech) and mRNA-1273 (Moderna), are based on the mRNA platform have shown promising results with efficacy above 90% (34, 35). Neither of these EUA COVID-19 vaccines are approved for use in pregnant women, breastfeeding mothers, children below the age of 18, and the immunocompromised (36), and they have yet to demonstrate long-lasting immune responses. Therefore, other vaccine approaches are still needed.

This study is the first to show the efficacy of a RABV vectored COVID-19 vaccine, CORAVAX, in the Syrian hamster model of severe disease. The rabies vaccine vector has several advantages: 1) it has an excellent safety profile, as it is used as an inactivated vaccine; 2) there is historical evidence of long-term immunity; 3) it can be administered safely and effectively to the vulnerable populations of children, pregnant women, elderly, and immunocompromised; 4) pre-existing rabies immunity does not affect the boosting potential of the vaccine; 5) RABV virions incorporate foreign antigens easily; and 6) RABV-based vaccines show excellent temperature stability (2, 19, 28).

Our vaccine approaches used only part of the SARS-CoV-2 spike protein. Using S1 rather than the full spike protein as the immunogen ensures the immune responses against the important neutralizing epitopes identified in human convalescent patients recognizing the RBD, NTD S1, and quaternary epitope that bridges the two RBDs (37). Our use of the S1 in CORAVAX might explain the higher VNA titers in the Syrian hamsters compared to other Covid-19 vaccines utilizing the full spike as the antigen in yellow fever and Ad26 vectors as well as in a DNA vaccine platform (38-40).

Several candidate vaccines against Covid-19 utilizing DNA, RNA, viral vectors, inactivated vaccines and subunit vaccine platforms are in various stages of pre-clinical or clinical development (41). While the immune correlates of protection are yet to be determined, most candidate vaccines trials compare their antibody and neutralizing response to SARS-CoV-2 convalescent sera. Clinical trials with convalescent plasma treatment have shown little or no significant difference in outcomes among SARS-CoV-2 patients compared to placebo group (42, 43). Also, most convalescent plasma samples obtained from individuals who recover from COVID-19 do not contain high levels of neutralizing activity (44).

CORAVAX induced high SARS CoV-2 S1 and RBD specific antibody and neutralizing titers on day 26 post the prime vaccination. The antibody and neutralizing titers were slightly (but not significantly) increased on day 56 after the boost (on day 28). CORAVAX also induced a strong Th1 biased immune response indicated by the IgG2/3 response before the challenge. Post-challenge, CORAVAX vaccinated animals induced significantly higher S1 IgG titers than the controls that correlated well with the neutralizing antibody response. Interestingly the RBD IgG titers were higher in the control animals at day 15 p.c. than CORAVAX vaccinating animals. The correlation of the S1 IgG titers with that of neutralizing antibody responses in the CORAVAX vaccinated animals suggests that the S1 IgG titers are a better predictor of protection than the RBD IgG titers. Our result aligns with the Chi et al. study that identified non-RBD binding, SARS CoV-2 S1 N terminal domain (NTD) binding neutralizing antibodies isolated from convalescent patients (45). It has also been suggested that the non-RBD binding S1 antibodies could be restraining the conformational changes of the S protein, thereby preventing viral entry. Conversely, vaccines that induce only RBD antibodies alone might induce resistance mutations in the virus (46). CORAVAX induces antibodies against the RBD as well other epitopes of S1. Antibodies induced in the CORAVAX vaccinated hamsters protected them from weight loss post challenge, while FILORAB1 vaccinated animals showed weight loss. The absence of infectious virus in the lungs and nasal turbinates of the CORAVAX vaccinated animals at day 3 p.c. suggests that CORAVAX can control viral transmission. Additionally, CORAVAX vaccinated animals showed significantly reduced lung pathology and inflammatory foci in comparison to the control group, suggesting that CORAVAX can dramatically reduce disease in vaccinated animals.

Regarding the adjuvant, we previously showed that GLA-SE adjuvanted FILORAB1 rabies virus-based EBOLA vaccine, protected 100% of NHPs challenged with Ebola, an improvement from the unadjuvanted FILORAB1 vaccine, which was less protective. This protection was attributed to the strong Th1 biased immune response induced by GLA-SE, which is a synthetic TRL4 agonist (20). To induce a stronger Th1 biased response in our study, we utilized another TRL4 agonist, MPLA-AddaVax, because MPLA is a TLR4 agonist and AddaVax is squalene-based oil-in-water nano-emulsion (similar to SE component in GLA-SE). MPLA has been shown to enhance the immunogenicity and protection of the rabies vaccine with induction of plasma cell responses. MPLA vaccinated mice accelerated the activation of dendritic cells, improving T-dependent B cell responses driving antibody production that skewed towards a strong Th1 bias (47). As seen for mild COVID-19 patients, a Th1 biased immune response is beneficial for protection protection against disease (48). Our previous work demonstrated that CORAVAX induced a strong Th1 biased immune response in Balb/C mice by generating higher IgG2a antibodies (30). In Syrian hamsters, CORAVAX vaccinated animals induced a stronger S1 IgG2/3 response before challenge. More importantly, the CORAVAX vaccinated hamsters mounted a significantly higher S1 IgG2/3 on day 15 p.c. than the FILORAB1 vaccinated animals. We could not detect S1 IgG1 responses, but we could detect RABV-G IgG1 responses, suggesting that S1 IgG1 antibodies are present at low levels or absent.

In conclusion, our RABV-vectored COVID-19 vaccine CORAVAX is efficacious and is able to prevent viral replication and reduce disease in Syrian hamsters. CORAVAX also serves as a vaccine against RABV because it induces high RABV VNA. Future studies assessing the efficacy of a single CORAVAX vaccine will be performed since we did not see a dramatic booster effect in immune responses among the hamsters. CORAVAX production should be relatively easy as it would follow the existing RABV vaccine manufacturing facilities and technologies. Additionally, the use of the adjuvant might allow for dose sparing. These results warrant further examination of CORAVAX in clinical trials to be conducted by Bharat Biotech International Ltd.

## MATERIALS AND METHODS

### Antibodies

The following SARS-CoV-2 specific human monoclonal antibodies were kindly provided by Distributed Bio: COVID-19 DB_A03-09, 12; DB_B01-04, B07-10, 12; DB_C01-05, 07, 09, 10; DB_D01, 02; DB_E01-04, 06, 07; DB_F02-03.

### Viruses

The SARS-CoV-2 strain used in this study is the first U.S. isolate SARS-CoV-2 USA_WA1/2020 from the Washington State patient identified on January 22, 2020 (31). Passage 3 was obtained from the World Reference Center for Emerging Viruses and Arboviruses (WRCEVA) at UTMB. Virus stocks were propagated in Vero E6 cells. The challenge stock used in this study is passage 5. The recombinant SARS-CoV-2 expressing Neon Green protein (SARS-CoV-2-mNG) (49) used in the neutralization assay was developed by Dr. Pei-Yong Shi at UTMB. Virus stocks were propagated in Vero E6 cells and a passage 4 was used.

### Vaccine production and purification

Recombinant RABV were recovered, purified, inactivated, and titered. Briefly, X-tremeGENE 9 transfection reagent (Millipore Sigma, Cat# 6365809001) was used to cotransfect the full-length viral cDNA clone encoding CORAVAX along with the plasmids encoding RABV N, P, and L proteins and the T7 RNA polymerase into BEAS-2B human lung cells in 6-well plates (RABV). At 8 days post transfection, supernatant was collected, filtered through a 0.22 µm membrane filter (Millipore), and titrated on VERO cells. The presence of recombinant virus was verified by immunostaining with monoclonal antibody against the nucleoprotein (FujiRebio, Cat# 800-092) and polyclonal antiserum against the S1 domain (Thermo Fisher, Cat# PA581798). The filtered virus was then used to inoculate VERO cells seeded in Cellstack Culture Chambers (Corning) and propagated in VP-SFM medium (Thermo Fisher Scientific) over a period of 18 days. Supernatant collected on day 10 post infection was filtered through 0.45 µm PES membrane filters (Nalgene) and layered onto 20% sucrose in DPBS. Virions were sedimented by ultracentrifugation in a SW32 rotor for 1.5 h at 25,000 rpm. Viral particles were resuspended in phosphate-buffered saline (PBS) and inactivated with 50 μl per mg of particles of a 1:100 dilution of β-propiolactone (BPL, Millipore Sigma, Cat# P5648) in cold water. The absence of infectious particles was verified by inoculating BSR cells with 10 μg of BPL-inactivated viruses over 3 passages.

### Animal studies

The studies were carried out in a strict accordance with the recommendations described in the Guide for the Care and Use of Laboratory Animals of the National Research Council. UTMB is an AAALAC-accredited institution and all animal work was approved by the IACUC Committee of UTMB. All efforts were made to minimize animal suffering and all procedures involving potential pain were performed with the appropriate anesthetic or analgesic. The number of hamsters used was scientifically justified based on statistical analyses of virological and immunological outcomes.

### Vaccination and SARS-CoV-2 challenge

Seven-week-old golden Syrian female hamsters (Envigo) were anesthetized with 5% isoflurane prior to immunization and blood collections and with ketamine/xylazine prior to the SARS-CoV-2 challenge. On day 0, 12 animals per group were inoculated with 10 μg of CORAVAX or FILORAB1 (control vaccine), adjuvanted with MPLA-AddaVax (Per animal: 5 μg MPLAs, InvivoGen, cat# vac-mpls; 50μL AddaVax, InvivoGen, cat#-adx-vac) in a 100 µl injection volume via the intramuscular route (50 µl per hind leg). The animals received a boost on day 28. Vena cava blood collections were performed two days prior the immunization, on day 26 and day 56. On day 60, vaccinated and control animals were exposed intranasally to the targeted dose of 10^5^ PFU of isolate SARS-CoV-2. Animals were monitored daily for weight loss and signs of disease. Half of the animals in each group (6 vaccinated and 6 control hamsters) was euthanized by overdose of injectable anesthetics on day 3 p.c. (day 63) for viral load determination. The remaining animals (5 vaccinated and 6 control hamsters) were euthanized on day 15 post challenge (p.c.) (day 75). One animal in the CORAVAX group did not recover from anesthesia after challenge.

### Recombinant proteins for ELISA

#### Purification of the HA-tagged protein from the supernatant of transfected cells for ELISA

Sub-confluent T175 flasks of 293T cells (human embryonic kidney cell line) were transfected with a pDisplay vector encoding amino acids 16 to 682 of SARS-CoV-2 S (S1) fused to a C-terminal hemagglutinin (HA) peptide using X-tremeGENE 9 reagent (Millipore Sigma, Cat# 6365809001). Supernatant was collected 6 days post-transfection, filtered through 0.22 um PES membrane filters (Nalgene) and the loaded onto an anti-HA agarose (Pierce, Cat# 26182) column equilibrated in PBS. After washing with ten bed volumes of PBS the column was loaded with 2 column volumes of HA peptide at a concentration of 400 µg/ml in PBS and incubated overnight at 4 °C. The following day, the protein was eluted with 2 column volumes of HA peptide followed by two column volumes of PBS. Fractions were collected and analyzed by western blotting with polyclonal antiserum against the S1 domain (Thermo Fisher, Cat# PA581798). Peak fractions were then pooled and dialyzed against PBS in 10,000 molecular weight cutoff (MWCO) dialysis cassettes (Thermo Fisher Scientific) to remove excess HA peptide. After dialysis, the protein was quantitated by UV spectrophotometry and frozen in small aliquots at −80 °C.

#### Purification of the RBD-His protein for ELISA

RBD-HIS (50): The SARS CoV-2 RBD-His tagged plasmid was purchased from Bei Resources (NR-52309). Sub-confluent T175 flasks of 293T cells (human embryonic kidney cell line) were transfected with the RBD-His tagged plasmid using X-tremeGENE 9 reagent (Millipore Sigma, Cat# 6365809001). Supernatant was collected 6 days post-transfection and filtered through 0.22 μm PES membrane filters (Nalgene). The 5 mL HisTALON cartridge (Clontech Laboratories, Cat # 635683) column was equilibrated with 10 column volumes of Equilibration Buffer (HisTALON™ Buffer Set, Clontech Laboratories, Cat# 635651). The filtered supernatant was loaded onto the HisTALON cartridge (Clontech Laboratories, Cat# 635683) column at a speed of 1 mL/min. After washing with ten column volumes of Wash Buffer (prepared by mixing 6.6 parts of Elution Buffer with 93.4 parts of Equilibration Buffer of the HisTALON™ Buffer Set), the sample is eluted (at a flow rate of ∼1 ml/min) with approximately 8 column volumes of Elution Buffer, collecting 1 ml fractions. The sample protein concentration was assessed by measuring the absorban 280 nm (Nanodrop, Thermo Fisher Scientific). Eluted fractions were analyzed by western blotting with a mouse monoclonal RBD specific antibody (InvivoGen, Cat# srbd-mab10). Peak fractions were then pooled and dialyzed against PBS in 10,000 molecular weight cutoff (MWCO) dialysis cassettes (Thermo Fisher Scientific). After dialysis, the protein was quantitated by UV spectrophotometry and frozen in small aliquots at −80 °C.

#### Enzyme-linked immunosorbent assay

To determine antibody responses to the S protein of SARS-CoV-2, an indirect ELISA was developed utilizing purified S1 or receptor binding domain (RBD) protein. The production of the recombinant proteins is described above. Humoral responses to SARS-CoV-2 S1 and RBD protein were measured by an indirect ELISA. We tested individual hamster sera by enzyme-linked immunosorbent assay (ELISA) for the presence of IgG specific to SARS-Cov-2 S1 or RBD. In order to test for anti-SARS CoV-2 S1 humoral responses, we produced soluble S1 or RBD as described above. The two recombinant proteins were resuspended in coating buffer (50 mM Na_2_CO_3_ [pH 9.6]) at a concentration of 0.5 μg/mL of S1 or 2 μg/mL of RBD, and then they were plated in 96-well ELISA MaxiSorp plates (Nunc) at 100 μl in each well. After overnight incubation at 4 °C, plates were washed 3 times with 1× PBST (0.05% Tween 20 in 1× PBS), followed by the addition of 250 μl blocking buffer (5% dry milk powder in 1× PBST) and incubation at room temperature for 1.5 h. The plates were then washed 3 times with PBST and incubated overnight at 4 °C with serial dilutions of sera (in triplicate) in 1× PBST containing 0.5% BSA. Plates were washed 3 times the next day, followed by the addition of HRP-conjugated goat anti-Syrian hamster IgG secondary antibody (Jackson immunoresearch, Cat# 107-035-142, 1:8000 in PBST) or mouse anti-hamster-IgG2/3-HRP (Southern Biotech, Cat# 1935-05, 1:8000 in PBST) or mouse anti-hamster-IgG1-HRP (Southern Biotech, Cat# 1940-05, 1:8000 in PBST) for 2 h at RT. After the incubation, plates were washed three times with PBST, and 200 μl of o-phenylenediamine dihydrochloride (OPD) substrate (Sigma) was added to each well. The reaction was stopped by the addition of 50 μl of 3 M H_2_SO_4_ per well. Optical density was determined at 490 nm (OD490) and 630 nm (OD630) using an ELX800 plate reader (Biotek Instruments, Inc., Winooski, VT). Plates were incubated for 15 min (IgG) or 20 min (IgG2/3 or IgG1) with OPD substrate before the reaction was stopped with 3 M H_2_SO_4_. Data were analyzed with GraphPad Prism (Version 8.0 g) using 4-parameter nonlinear regression to determine the titer at which the curves reach 50% of the top plateau value (50% effective concentration [EC50]).

#### Neutralizing antibody response

Sera collected from animals were tested for neutralizing capabilities against SARS-CoV-2. Briefly, serum samples were heat-inactivated (30 min at 56°C), and then 10-fold diluted sera were further diluted in a 2-fold *serial* fashion, and 60 µl of each serum dilution was mixed with 60 µl of SARS-CoV-2-mNG (200 PFU) (49). The serum/virus mixtures were incubated for 1 h at 37°C. 100 µl of the serum/virus mixtures were then transferred to Vero E6 cell monolayers in flat-bottom 96-well plates and incubated for 2 days at 37°C. Virus fluorescence was measured with a Cytation Hybrid Multi-Mode reader at 488 nm (Biotek Instruments).

#### Tissue processing and viral load determination

Animals were euthanized on days 3 and 15 p.c., and lungs and nasal turbinates were harvested. Right lungs and nasal turbinates were placed in L15 medium supplemented with 10% fetal bovine serum (Gibco) and Antibiotic-Antimycotic solution (Gibco). Tissues were homogenized using the TissueLyser II system (Qiagen) and tissue homogenates were aliquoted and stored at -80°C. Tissue homogenates were titrated on Vero E6 cell monolayers in 24-well plates to determine viral loads. Plates were incubated 3 days at 37°C before being fixed with 10% formalin and removed from containment. Plaques were visualized by immunostaining with 1 µg/mL cocktail of 37 SARS-CoV-2 specific human antibodies kindly provided by Distributed Bio. As the secondary antibody, HRP-labeled goat anti-human IgG (SeraCare) was used at dilution 1:500. Primary and secondary antibodies were diluted in 1X DPBS with 5% milk. Plaques were revealed by AEC substrate (enQuire Bioreagents).

#### Viral RNA copies by qRT-PCR

Tissue homogenates were mixed with TRIzol Reagent (Life Technologies) at a 1:5 volume ratio of homogenate to TRIzol. The RNA extraction protocol for biological fluids using TRIzol Reagent was followed until the phase separation step. The remaining RNA extraction was done using the PureLink RNA Mini Kit (Ambion). The quantity and quality (260/280 ratios) of RNA extracted was measured using NanoDrop (Thermo Fisher). SARS-CoV-2 nucleoprotein cDNA was generated from RNA from Bei Resources (NR-52285) by One-Step RT PCR (SuperScript IV, Thermo Fisher) with primers SARS CoV-2 N IVT F1 (5’-GAATTCTAATACGACTCACTATAGGGGATGTCTGATAATGGACCC-3’) and SARS CoV-2 N IVT R1 (5’-GCTAGCTTAGGCCTGAGTTGAGTCAGCACTGCT-3’). The SARS-CoV-2 N standards were generated by *in-vitro* transcription of the generated SARS-CoV-2 N cDNA using the MegaScript T7 Transcription kit (Invitrogen), followed by using the MEGAclear Transcription Clean-Up Kit. Aliquots of 2 ×10^10^ copies/µL were frozen at -80°C. Five microliters of RNA per sample were run in triplicate, using the primers 2019-nCoV_N2-F (5’-TTACAAACATTGGCCGCAAA-3’), 2019-nCoV_N2-R (5’-GCGCGACATTCCGAAGAA-3’) and probe 2019-nCoV_N2-P-FAM (5’-ACAATTTGCCCCCAGCGCTTCAG-3’).

#### Histopathology

Following euthanasia, necropsy was performed, gross lesions were noted, and left lungs were placed in 10% formalin for histopathological assessment. After a 24-h incubation at 4°C, lungs were transferred to fresh 10% formalin for an additional 48-hour incubation before removal from containment. Tissues were processed by standard histological procedures by the UTMB Anatomic Pathology Core, and 4 μm-thick sections were cut and stained with hematoxylin and eosin. Sections of lungs were examined for the extent of inflammation, type of inflammatory foci, and changes in alveoli/alveolar septa/airways/blood vessels in parallel with sections from uninfected or unvaccinated lungs. The blinded tissue sections were semi-quantitatively scored for pathological lesions using the criteria described in Supplemental Table 1. Examination was performed with an Olympus CX43 microscope at magnification 40X for general observation and 100X magnification for detailed observation. Each section was scored by a trained member of staff.

#### Statistical analysis

Statistical analyses were performed with GraphPad Prism for Windows (version 6.07). *P*<0.05 was considered significant. P > 0.123 (ns), P < 0.033 (*), P < 0.002 (**), P < 0.001 (***).

#### Biocontainment work

Work with SARS-CoV-2 was performed in the BSL-4 facilities of the Galveston National Laboratory according to approved standard operating protocols.

## Acknowledgments

This work was supported in part by a grant from Bharat Biotech, India, and the State of Pennsylvania (add number) to MJS and the Jefferson Vaccine Center. We thank Jennifer Wilson (Thomas Jefferson University, Philadelphia, PA) for critical reading and editing of the manuscript.

## Author Contributions

D.K performed the ELISA, RNA isolations, and qRT-PCR assays and wrote the manuscript. D.C.M. designed and supervised the animal study, isolated tissues, performed tissue viral load determination, neutralization assays. C.W. designed and produced the vaccine. R.L. isolated RNA and performed the RFFIT assay. A.J.R. analyzed the pathology slides. L.Z.D. performed ELISAs. M.J.S. and A.B. conceived and supervised the study, secured funding and edited the manuscript.

## Supplemental Information

**Supplemental Table 1: Histopathology scoring criteria**

## References

1. Anonymous. World Health Organization.

2. Anonymous. Centers for Disease Control and Prevention.

3. Kaiser J. Temperature concerns could slow the rollout of new coronavirus vaccines. https://www.sciencemag.org/news/2020/11/temperature-concerns-could-slow-rollout-new-coronavirus-vaccines.

4. Leitner WW, Ying H, Restifo NP. 1999. DNA and RNA-based vaccines: principles, progress and prospects. Vaccine 18:765–77.

5. Garg H, Mehmetoglu-Gurbuz T, Joshi A. 2020. Virus Like Particles (VLP) as multivalent vaccine candidate against Chikungunya, Japanese Encephalitis, Yellow Fever and Zika Virus. Sci Rep 10:4017.

6. Roldao A, Mellado MC, Castilho LR, Carrondo MJ, Alves PM. 2010. Virus-like particles in vaccine development. Expert Rev Vaccines 9:1149–76.

7. Anonymous. GAVI. https://www.gavi.org/vaccineswork/what-are-viral-vector-based-vaccines-and-how-could-they-be-used-against-covid-19.

8. Ura T, Okuda K, Shimada M. 2014. Developments in Viral Vector-Based Vaccines. Vaccines (Basel) 2:624–41.

9. Lauer KB, Borrow R, Blanchard TJ. 2017. Multivalent and Multipathogen Viral Vector Vaccines. Clin Vaccine Immunol 24.

10. Papaneri AB, Wirblich C, Marissen WE, Schnell MJ. 2013. Alanine scanning of the rabies virus glycoprotein antigenic site III using recombinant rabies virus: implication for postexposure treatment. Vaccine 31:5897–902.

11. Blaney JE, Marzi A, Willet M, Papaneri AB, Wirblich C, Feldmann F, Holbrook M, Jahrling P, Feldmann H, Schnell MJ. 2013. Antibody quality and protection from lethal Ebola virus challenge in nonhuman primates immunized with rabies virus based bivalent vaccine. PLoS Pathog 9:e1003389.

12. Schnell MJ, Tan GS, Dietzschold B. 2005. The application of reverse genetics technology in the study of rabies virus (RV) pathogenesis and for the development of novel RV vaccines. J Neurovirol 11:76–81.

13. Schnell MJ, McGettigan JP, Wirblich C, Papaneri A. 2010. The cell biology of rabies virus: using stealth to reach the brain. Nat Rev Microbiol 8:51–61.

14. Gomme EA, Faul EJ, Flomenberg P, McGettigan JP, Schnell MJ. 2010. Characterization of a single-cycle rabies virus-based vaccine vector. J Virol 84:2820–31.

15. Lawrence TM, Wanjalla CN, Gomme EA, Wirblich C, Gatt A, Carnero E, Garcia-Sastre A, Lyles DS, McGettigan JP, Schnell MJ. 2013. Comparison of Heterologous Prime-Boost Strategies against Human Immunodeficiency Virus Type 1 Gag Using Negative Stranded RNA Viruses. PLoS One 8:e67123.

16. McGettigan JP, Koser ML, McKenna PM, Smith ME, Marvin JM, Eisenlohr LC, Dietzschold B, Schnell MJ. 2006. Enhanced humoral HIV-1-specific immune responses generated from recombinant rhabdoviral-based vaccine vectors co-expressing HIV-1 proteins and IL-2. Virology 344:363–77.

17. Walsh PD, Kurup D, Hasselschwert DL, Wirblich C, Goetzmann JE, Schnell MJ. 2017. The Final (Oral Ebola) Vaccine Trial on Captive Chimpanzees? Sci Rep 7:43339.

18. McKenna PM, Koser ML, Carlson KR, Montefiori DC, Letvin NL, Papaneri AB, Pomerantz RJ, Dietzschold B, Silvera P, McGettigan JP, Schnell MJ. 2007. Highly attenuated rabies virus-based vaccine vectors expressing simian-human immunodeficiency virus89.6P Env and simian immunodeficiency virusmac239 Gag are safe in rhesus macaques and protect from an AIDS-like disease. J Infect Dis 195:980–8.

19. Kurup D, Fisher CR, Smith TG, Abreu-Mota T, Yang Y, Jackson FR, Gallardo-Romero N, Franka R, Bronshtein V, Schnell MJ. 2019. Inactivated Rabies Virus-Based Ebola Vaccine Preserved by Vaporization Is Heat-Stable and Immunogenic Against Ebola and Protects Against Rabies Challenge. J Infect Dis 220:1521–1528.

20. Johnson RF, Kurup D, Hagen KR, Fisher C, Keshwara R, Papaneri A, Perry DL, Cooper K, Jahrling PB, Wang JT, Ter Meulen J, Wirblich C, Schnell MJ. 2016. An Inactivated Rabies Virus-Based Ebola Vaccine, FILORAB1, Adjuvanted With Glucopyranosyl Lipid A in Stable Emulsion Confers Complete Protection in Nonhuman Primate Challenge Models. J Infect Dis 214:S342–S354.

21. Fisher CR, Schnell MJ. 2018. New developments in rabies vaccination. Rev Sci Tech 37:657–672.

22. Abreu-Mota T, Hagen KR, Cooper K, Jahrling PB, Tan G, Wirblich C, Johnson RF, Schnell MJ. 2018. Non-neutralizing antibodies elicited by recombinant Lassa-Rabies vaccine are critical for protection against Lassa fever. Nat Commun 9:4223.

23. Wirblich C, Coleman CM, Kurup D, Abraham TS, Bernbaum JG, Jahrling PB, Hensley LE, Johnson RF, Frieman MB, Schnell MJ. 2017. One-Health: a Safe, Efficient, Dual-Use Vaccine for Humans and Animals against Middle East Respiratory Syndrome Coronavirus and Rabies Virus. J Virol 91.

24. Keshwara R, Shiels T, Postnikova E, Kurup D, Wirblich C, Johnson RF, Schnell MJ. 2019. Rabies-based vaccine induces potent immune responses against Nipah virus. NPJ Vaccines 4:15.

25. Keshwara R, Hagen KR, Abreu-Mota T, Papaneri AB, Liu D, Wirblich C, Johnson RF, Schnell MJ. 2019. A Recombinant Rabies Virus Expressing the Marburg Virus Glycoprotein Is Dependent upon Antibody-Mediated Cellular Cytotoxicity for Protection against Marburg Virus Disease in a Murine Model. J Virol 93.

26. Hudacek AW, Al-Saleem FH, Willet M, Eisemann T, Mattis JA, Simpson LL, Schnell MJ. 2014. Recombinant rabies virus particles presenting botulinum neurotoxin antigens elicit a protective humoral response in vivo. Mol Ther Methods Clin Dev 1:14046.

27. Kurup D, Wirblich C, Feldmann H, Marzi A, Schnell MJ. 2015. Rhabdovirus-based vaccine platforms against henipaviruses. J Virol 89:144–54.

28. Scher G, Schnell MJ. 2020. Rhabdoviruses as vectors for vaccines and therapeutics. Curr Opin Virol 44:169–182.

29. Fisher CR, Streicker DG, Schnell MJ. 2018. The spread and evolution of rabies virus: conquering new frontiers. Nat Rev Microbiol 16:241–255.

30. Kurup D, Wirblich C, Ramage H, Schnell MJ. 2020. Rabies virus-based COVID-19 vaccine CORAVAX induces high levels of neutralizing antibodies against SARS-CoV-2. NPJ Vaccines 5:98.

31. Harcourt J, Tamin A, Lu X, Kamili S, Sakthivel SK, Murray J, Queen K, Tao Y, Paden CR, Zhang J, Li Y, Uehara A, Wang H, Goldsmith C, Bullock HA, Wang L, Whitaker B, Lynch B, Gautam R, Schindewolf C, Lokugamage KG, Scharton D, Plante JA, Mirchandani D, Widen SG, Narayanan K, Makino S, Ksiazek TG, Plante KS, Weaver SC, Lindstrom S, Tong S, Menachery VD, Thornburg NJ. 2020. Severe Acute Respiratory Syndrome Coronavirus 2 from Patient with Coronavirus Disease, United States. Emerg Infect Dis 26:1266–1273.

32. Sia SF, Yan LM, Chin AWH, Fung K, Choy KT, Wong AYL, Kaewpreedee P, Perera R, Poon LLM, Nicholls JM, Peiris M, Yen HL. 2020. Pathogenesis and transmission of SARS-CoV-2 in golden hamsters. Nature 583:834–838.

33. Cevik M, Kuppalli K, Kindrachuk J, Peiris M. 2020. Virology, transmission, and pathogenesis of SARS-CoV-2. BMJ 371:m3862.

34. Polack FP, Thomas SJ, Kitchin N, Absalon J, Gurtman A, Lockhart S, Perez JL, Perez Marc G, Moreira ED, Zerbini C, Bailey R, Swanson KA, Roychoudhury S, Koury K, Li P, Kalina WV, Cooper D, Frenck RW, Jr., Hammitt LL, Tureci O, Nell H, Schaefer A, Unal S, Tresnan DB, Mather S, Dormitzer PR, Sahin U, Jansen KU, Gruber WC, Group CCT. 2020. Safety and Efficacy of the BNT162b2 mRNA Covid-19 Vaccine. N Engl J Med 383:2603–2615.

35. Anonymous. November 16, 2020. Promising Interim Results from Clinical Trial of NIH-Moderna COVID-19 Vaccine. NIAID Office of Communications. https://www.nih.gov/news-events/news-releases/promising-interim-results-clinicaltrial-nih-moderna-covid-19-vaccine.

36. Anonymous. U.S. Food and Drug Administration.

37. Liu L, Wang P, Nair MS, Yu J, Rapp M, Wang Q, Luo Y, Chan JF, Sahi V, Figueroa A, Guo XV, Cerutti G, Bimela J, Gorman J, Zhou T, Chen Z, Yuen KY, Kwong PD, Sodroski JG, Yin MT, Sheng Z, Huang Y, Shapiro L, Ho DD. 2020. Potent neutralizing antibodies against multiple epitopes on SARS-CoV-2 spike. Nature 584:450–456.

38. Tostanoski LH, Wegmann F, Martinot AJ, Loos C, McMahan K, Mercado NB, Yu J, Chan CN, Bondoc S, Starke CE, Nekorchuk M, Busman-Sahay K, Piedra-Mora C, Wrijil LM, Ducat S, Custers J, Atyeo C, Fischinger S, Burke JS, Feldman J, Hauser BM, Caradonna TM, Bondzie EA, Dagotto G, Gebre MS, Jacob-Dolan C, Lin Z, Mahrokhian SH, Nampanya F, Nityanandam R, Pessaint L, Porto M, Ali V, Benetiene D, Tevi K, Andersen H, Lewis MG, Schmidt AG, Lauffenburger DA, Alter G, Estes JD, Schuitemaker H, Zahn R, Barouch DH. 2020. Ad26 vaccine protects against SARS-CoV-2 severe clinical disease in hamsters. Nat Med 26:1694–1700.

39. Sanchez-Felipe L, Vercruysse T, Sharma S, Ma J, Lemmens V, Van Looveren D, Javarappa MPA, Boudewijns R, Malengier-Devlies B, Liesenborghs L, Kaptein SJF, De Keyzer C, Bervoets L, Debaveye S, Rasulova M, Seldeslachts L, Li LH, Jansen S, Yakass MB, Verstrepen BE, Boszormenyi KP, Kiemenyi-Kayere G, van Driel N, Quaye O, Zhang X, Ter Horst S, Mishra N, Deboutte W, Matthijnssens J, Coelmont L, Vandermeulen C, Heylen E, Vergote V, Schols D, Wang Z, Bogers W, Kuiken T, Verschoor E, Cawthorne C, Van Laere K, Opdenakker G, Velde GV, Weynand B, Teuwen DE, Matthys P, Neyts J, Jan Thibaut H, Dallmeier K. 2020. A single-dose live-attenuated YF17D-vectored SARS-CoV-2 vaccine candidate. Nature doi:10.1038/s41586-020-3035-9.

40. Rebecca L. Brocato SAK, Robert K. Kim, Xiankun Zeng, Lucia M. Principe, Jeffrey M. Smith, Jay W. Hooper. 2020. Protective efficacy of a SARS-CoV-2 DNA Vaccine in wild-type and immunosuppressed Syrian hamsters. bioRxiv 2020.11.10.376905; doi: https://doi.org/10.1101/2020.11.10.37690.

41. Dong Y, Dai T, Wei Y, Zhang L, Zheng M, Zhou F. 2020. A systematic review of SARS-CoV-2 vaccine candidates. Signal Transduct Target Ther 5:237.

42. Simonovich VA, Burgos Pratx LD, Scibona P, Beruto MV, Vallone MG, Vazquez C, Savoy N, Giunta DH, Perez LG, Sanchez MDL, Gamarnik AV, Ojeda DS, Santoro DM, Camino PJ, Antelo S, Rainero K, Vidiella GP, Miyazaki EA, Cornistein W, Trabadelo OA, Ross FM, Spotti M, Funtowicz G, Scordo WE, Losso MH, Ferniot I, Pardo PE, Rodriguez E, Rucci P, Pasquali J, Fuentes NA, Esperatti M, Speroni GA, Nannini EC, Matteaccio A, Michelangelo HG, Follmann D, Lane HC, Belloso WH, PlasmAr Study G. 2020. A Randomized Trial of Convalescent Plasma in Covid-19 Severe Pneumonia. N Engl J Med doi:10.1056/NEJMoa2031304.

43. Liu STH, Lin HM, Baine I, Wajnberg A, Gumprecht JP, Rahman F, Rodriguez D, Tandon P, Bassily-Marcus A, Bander J, Sanky C, Dupper A, Zheng A, Nguyen FT, Amanat F, Stadlbauer D, Altman DR, Chen BK, Krammer F, Mendu DR, Firpo-Betancourt A, Levin MA, Bagiella E, Casadevall A, Cordon-Cardo C, Jhang JS, Arinsburg SA, Reich DL, Aberg JA, Bouvier NM. 2020. Convalescent plasma treatment of severe COVID-19: a propensity score-matched control study. Nat Med 26:1708–1713.

44. Robbiani DF, Gaebler C, Muecksch F, Lorenzi JCC, Wang Z, Cho A, Agudelo M, Barnes CO, Gazumyan A, Finkin S, Hagglof T, Oliveira TY, Viant C, Hurley A, Hoffmann HH, Millard KG, Kost RG, Cipolla M, Gordon K, Bianchini F, Chen ST, Ramos V, Patel R, Dizon J, Shimeliovich I, Mendoza P, Hartweger H, Nogueira L, Pack M, Horowitz J, Schmidt F, Weisblum Y, Michailidis E, Ashbrook AW, Waltari E, Pak JE, Huey-Tubman KE, Koranda N, Hoffman PR, West AP, Jr., Rice CM, Hatziioannou T, Bjorkman PJ, Bieniasz PD, Caskey M, Nussenzweig MC. 2020. Convergent antibody responses to SARS-CoV-2 in convalescent individuals. Nature 584:437–442.

45. Chi X, Yan R, Zhang J, Zhang G, Zhang Y, Hao M, Zhang Z, Fan P, Dong Y, Yang Y, Chen Z, Guo Y, Zhang J, Li Y, Song X, Chen Y, Xia L, Fu L, Hou L, Xu J, Yu C, Li J, Zhou Q, Chen W. 2020. A neutralizing human antibody binds to the N-terminal domain of the Spike protein of SARS-CoV-2. Science 369:650–655.

46. Zhou H, Chen Y, Zhang S, Niu P, Qin K, Jia W, Huang B, Zhang S, Lan J, Zhang L, Tan W, Wang X. 2019. Structural definition of a neutralization epitope on the N-terminal domain of MERS-CoV spike glycoprotein. Nat Commun 10:3068.

47. Chen C, Zhang C, Li R, Wang Z, Yuan Y, Li H, Fu Z, Zhou M, Zhao L. 2019. Monophosphoryl-Lipid A (MPLA) is an Efficacious Adjuvant for Inactivated Rabies Vaccines. Viruses 11.

48. Allegra A, Di Gioacchino M, Tonacci A, Musolino C, Gangemi S. 2020. Immunopathology of SARS-CoV-2 Infection: Immune Cells and Mediators, Prognostic Factors, and Immune-Therapeutic Implications. Int J Mol Sci 21.

49. Xie X, Muruato A, Lokugamage KG, Narayanan K, Zhang X, Zou J, Liu J, Schindewolf C, Bopp NE, Aguilar PV, Plante KS, Weaver SC, Makino S, LeDuc JW, Menachery VD, Shi PY. 2020. An Infectious cDNA Clone of SARS-CoV-2. Cell Host Microbe 27:841–848 e3.

50. Stadlbauer D, Amanat F, Chromikova V, Jiang K, Strohmeier S, Arunkumar GA, Tan J, Bhavsar D, Capuano C, Kirkpatrick E, Meade P, Brito RN, Teo C, McMahon M, Simon V, Krammer F. 2020. SARS-CoV-2 Seroconversion in Humans: A Detailed Protocol for a Serological Assay, Antigen Production, and Test Setup. Curr Protoc Microbiol 57:e100.

